# Insight into the Thermo-responsive Phase Behavior of P1 Domain of *α*-synuclein using Atomistic Simulations

**DOI:** 10.1101/2024.12.14.628487

**Authors:** Sanchari Chakraborty, Mithun Biswas

## Abstract

Biomolecular condensate formation driven by intrinsically disordered proteins (IDPs) is regulated by interaction between various domains of the protein. Such condensates are implicated in various neurodegenerative diseases. The presynaptic intrinsically disordered protein, *α*-Syn is involved in the pathogenesis of Parkinson’s disease. The central non-amyloid *β*-component (NAC) domain in the protein is considered to be a major driver of pathogenic aggregation, although recent studies have suggested that the P1 domain from the flanking N-terminal region, can act as a ‘master controller’ for *α*-Syn function and aggregation. To gain molecular insight into the phase behavior of the P1 domain itself, we investigate how assemblies of P1 (residues 36-42) chains phase separate with varying temperatures using all-atom molecular dynamics simulations. The simulations reveal that P1 is able to phase separate above a lower critical solution temperature. Formation of the condensed phase is driven by exclusion of water molecules by the hydrophobic chains. P1 chain density in the condensate is determined by weak multi-chain interactions between the residues. Moreover, Tyrosine (Y^39^) is involved in the formation of strongest contacts between residue pairs in the dense phase. These results provide a detailed picture of condensate formation by a key segment of the *α*-Syn molecule, which may help deciphering the phase behavior of the full-length protein.

## 1 Introduction

The crowded intracellular medium promotes interactions between ordered and disordered proteins,^1,2^ oligomerization,^3,4^ signal transduction^5^ and enzymatic activity.^6,7^ Recent evidences suggest that a part of this medium is formed as membraneless assemblies of protein, RNA and other biomolecules exhibiting liquid-like properties which can also perform specific tasks.^8–16^ Intrinsically disordered proteins (IDPs), such as *α*-Syn, TDP-43, FUS, or peptides having intrinsically disordered regions (IDRs), facilitate formation of these membraneless assemblies by stabilizing condensed liquid-like phases in a process called liquid-liquid phase separation (LLPS).^13,15,17,18^ Recently, many IDPs have been reported to undergo LLPS prior to formation of amyloid fibrils linked to diseases like Parkinson’s,^19,20^ amyotrophic lateral sclerosis (ALS),^21^ frontotemporal dementia^22^ and Alzheimer’s.^23^ From the evidences it seems likely that increase in the concentration of IDPs inside the phase-separated compartments may aid formation of pathological aggregates.^24^

Weak multivalent interactions between IDPs are major contributors towards LLPS.^15,16,25,26^ The presence of low complexity domains (LCDs) and lack of stable secondary structure make IDPs highly susceptible to LLPS. Environmental factors such as pH, salt concentration, temperature, presence of membranes, and crowding elements (osmolytes) influence the formation of condensates as well.^27,28^ IDPs exhibit distinct response to thermal stimulus depending on their sequence composition such as upper critical solution temperature (UCST), lower critical solution temperature(LCST), or both UCST and LCST.^18,29,30^

*α*-Syn is a well-studied 140 residue long IDP present within the presynapse of human brain which undergoes LLPS prior to aggregation.^20^ Altered dynamics of *α*-Syn leads to the formation of pathogenic aggregates involved in Parkinson’s disease, multiple system atrophy, and dementia.^31,32^ The three primary structural domains of *α*-Syn, N-terminal (residues 1-60), the non-amyloid *β* component (NAC) (residues 61-95), and the C-terminal region (residues 95-140), are predicted to have distinct roles in forming higher order assembly by the protein.^32,33^ The N-terminal is comprised of KTKEGV amino acid repeat motifs and can bind to membranes.^16,34^ Most of the familial PD related mutants such as A30P, E46K, and A53T occur in this region.^34^ The NAC domain is rich in hydrophobic residues and is the major driver of pathogenic aggregation. The C-terminal tail is highly disordered, acidic and can interact with metal ions and small molecules. It has been predicted that long range contacts between the N-and C-terminals may regulate the accessibility of the NAC region to form intermolecular interactions.^35^ Despite the crucial part NAC plays in *α*-Syn function and aggregation, increasing evidences depict the importance of its flanking domains in propagating pathogenic aggregation.^32,36,37^

Recently, two fragments, namely P1 (residue 36-42) and P2 (residue 45-57), within the N-terminal have been found to be essential for *α*-Syn aggregation.^38^ Deletion of both the P1 and P2 sequences inhibits the protein’s ability to aggregate. P1 is termed as a master controller of *α*-Syn function such that replacing it with glycine-serine repeats inhibits aggregation suggesting high sequence specificity.^34^ P1 region is centered on residue Y^39^ which binds various chaperons and is also considered to be a modulator of the peptide structure and aggregation kinetics.^34,39–41^ A peptide sequence comprising both P1 and P2 (residue 36-55) readily self assemble in vitro^42^ and in silico^43^ and display formation of *β*-hairpins that self-assemble into oligomers.^42^

Although significant efforts have been made through microscopy, fluorescence techniques, solution-state NMR and particle-tracking microrheology to study LLPS in recent years,^15,44–46^ understanding how residue-level interactions regulate LLPS behavior remains a challenging task. All-atom molecular dynamics (MD) simulations are a promising alternative to explore the phenomenon at a molecular level. Historically, allatom models have been used to study globular proteins, but recently they are being adapted for IDPs as well.^47^ As the computational cost to capture phase separation of IDPs through all-atom simulations is very high, few studies to date that have been reported do not capture the phase behavior through direct simulations.^48–52^ However, direct coexistence MD simulations of short peptides have been employed previously to study the LLPS behavior.^53^ To overcome the limitations of all-atom models, coarse grained models, such as Martini,^54^ which maps four atoms to one CG bead, has been employed to study phase separation of proteins.^55–58^ A further simplified CG strategy, the HPS model,^59^ where each residue in a protein is represented by a single bead, combined with the slab coexistence simulation approach has also been successful in determining the dense and dilute protein concentrations and critical temperatures for large IDP assemblies.^60,61^ The patchy-colloid models, in which the entire protein is represented by a sphere with interaction sites or patches on them, have been used to examine the effect of valence of proteins, and also to study protein-RNA condensation mechanisms.^62–65^ All of these studies greatly improve our understanding of the phase separation phenomenon of IDPs. However, simplified CG models lack a detailed picture of residue-residue and residue-water interactions that help stabilize condensate formation.

Here, we explore the condensed phase formation of P1 domain of *α*-Syn employing atomistic MD simulation. Although several attempts, including fragment based approaches, have been made in recent years to study *α*-Syn oligomers and fibrillation using MD simulations,^43,66–71^ this is the first study to the best of our knowledge, to investigate LLPS behavior of a fragment peptide (the crucial P1 domain of *α*-Syn) through direct coexistence simulation. As the experimental evidence suggests that P1 fragment acts as a master regulator of *α*-Syn aggregation, it seems likely that it may play a key role in the phase separation of the protein as well. Our results demonstrate that multiple copies of P1 domains can self assemble and phase separate exhibiting a lower critical solution temperature. We then investigate and elaborate upon the underlying network of interactions driving its phase behavior.

## 2 Methods

### 2.1 System Preparation and MD Simulations

To explore the phase separation characteristics, allatom molecular dynamics simulations of the P1 region (^36^GVLYVGS^42^) of *α*-Syn (PDB ID-1XQ8) (see ESI Fig.S1)† were carried out employing Amber ff99SB-disp force field with modified TIP4P-D water model.^72^ For comparison with previous coexistence simulation of peptides,^53^ independent simulations employing Amber ff99SB-ILDN force field^73^ with TIP3P water model were also performed (see ESI Table S1)†. The N and C termini of the P1 peptide were capped with ACE and NME groups respectively, using Pymol.^74^ Thereafter, 70 copies of P1 chains were inserted into a box of dimensions 6×6×9 nm^3^ with PACKMOL.^75^ Variation in the chain density profile was not observed for a box with longer 𝒵-dimension employing Amber ff99SB-ILDN force field (see ESI Fig. S2)†. The system was then solvated explicitly by adding appropriate number of water molecules, resulting in a concentration of 0.35M for peptide chains. The system is first energy minimized using steepest descent for 50000 steps. Then NVT equilibration was performed for 2 ns with position restraints on the heavy protein atoms at a temperature of 300 K. This was followed by another equilibration simulation under NPT conditions for 2 ns using Parinello-Rahman barostat with isotropic pressure coupling at a reference pressure of 1 bar.

In order to investigate the thermoresponsive phase behavior, 15 temperatures were chosen between 265-350 K (see ESI Table S1†) and at each temperature 600 ns long NVT production run was carried out using GROMACS.^76^ Formation of condensed phase was observed for all the simulations employing Amber ff99SB-ILDN force field, but simulations with Amber ff99SB-disp force field failed to phase separate below 293 K. For both force fields, it was observed that the formation of protein-dense phase requires approximately 300 ns (detailed later). Thus the equilibrium properties of the coexisting phases were estimated from the last 300 ns of each simulation. At each temperature, the dense and dilute phases have been identified using a cutoff-based approach (see ESI Fig.S2)†.

### 2.2 Determination of critical temperature

The critical temperature *T*_*C*_ is obtained by fitting dense and dilute phase densities, *ρ*_dense_ and *ρ*_dilute_ respectively, to the following equation

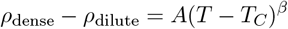

where, *β*=0.325 is the critical exponent (for 3D Ising universality class) and *A* is a fitting constant. To choose an optimal range for fitting temperatures for determination of *T*_*C*_, we choose a fixed maximum temperature (*T*_*max*_) and calculate the relative error in fitting for various values of minimum temperature (*T*_*min*_). *T*_*C*_ corresponds to the temperature range with minimum error value. Variation of *T*_*C*_ for different choices of *T*_*min*_ and *T*_*max*_ is given in the ESI (Table S2 and Table S3)†

### 2.3 Trajectory analysis

The simulation trajectories were analyzed using various tools available within the GROMACS package. Molecular trajectories were viewed using VMD.^77^ The error bars have been estimated by block averaging over the final 300 ns of the simulations.

## 3 Results and Discussion

### 3.1 Condensate formation in P1-water binary mixture

A set of simulations were carried out consisting of multiple copies of capped P1 (^36^GVLYVGS^42^) domain at different temperatures. For simulations with Amber ff99SB-disp force field, the peptides formed condensates above 293 K, but for Amber ff99SB-ILDN force field, the dispersed peptide chains came together forming a dense slab like configuration during the entire temperature range 265-350 K. Nevertheless, the chains remain highly dynamic throughout the simulation and fail to hold any particular conformation for a longer duration.

To further investigate the condensate formation, we calculated the solvent accessible surface area (SASA) of the peptide chains as a function of simulation time at two different temperatures, at 290 and 350 K, (Fig. 1a). Employing Amber ff99SB-disp force field at 350 K (*yellow trace*), the SASA value decreases from its initial value and stabilize approximately after 300 ns. However, at 290 K (*red trace*) P1 chains do not form a condensed phase during the entire 600 ns simulation as revealed by continuously decreasing SASA values. Simulations with the Amber ff99SB-ILDN force field also stabilize after first 300 ns of the simulation and shows little change during the last 300 ns indicating formation of stable P1 chain clusters (data not shown). Hence, all subsequent analysis for both the force fields was performed on the final 300 ns of the simulation trajectories. Two metrics have been employed to characterize condensate formation by the peptide chains, namely the aggregation propensity (AP) and clustering degree (CD).^53,56^ The aggregation propensity of protein chains is described as the ratio of the initial SASA to the instantaneous SASA. Fig. 1b plots the average AP values (over the last 300 ns) at various temperatures showing that AP values increase monotonically with temperature. Clustering degree captures formation of large condensates and is defined as the ratio of number of protein molecules in the largest cluster to the total number of proteins in the box. Here, two chains are considered as part of the same cluster if the distance between atoms in two chains is less than 0.2 nm. Similar to AP, variation of CD values as function of temperature shows that more protein chains become part of the largest cluster as temperature increases upto 320 K, when only a few chains remain dispersed outside the dense slab.

**Figure 1:**
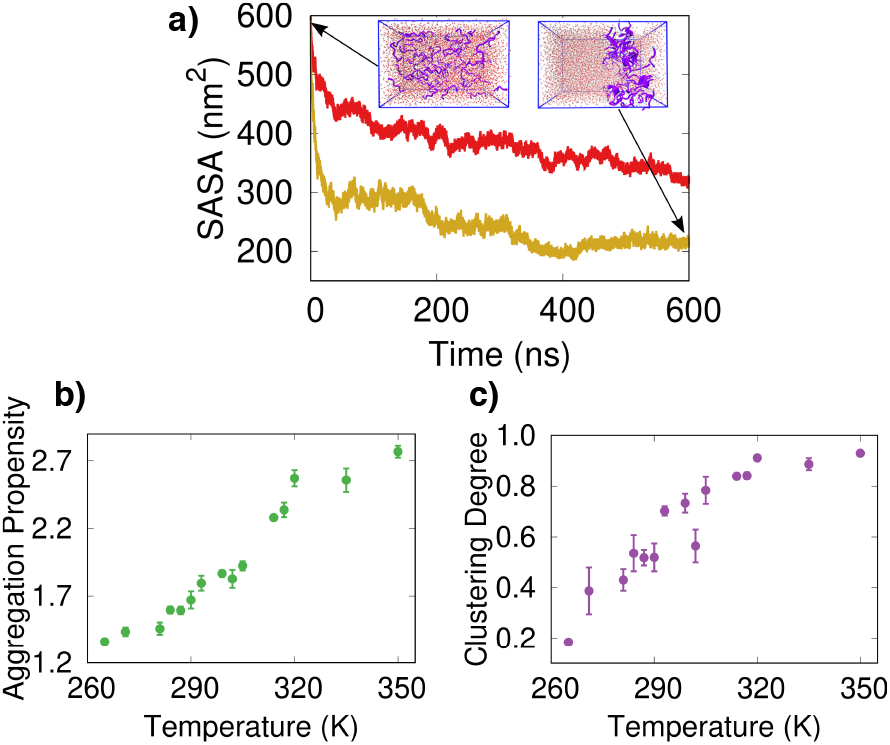
Condensate formation by P1 chains using Amber ff99SB-disp force field. (a) Time evolution of SASA values for assembly of P1 chains at 290 (*red* trace) and 350 K (*yellow* trace). (b)-(c) AP and CD values for P1 chains as function of temperature.

It is known that many all-atom force-fields well-suited for globular protein simulations perform poorly when applied to disordered peptides.^78–81^ Standard MD force-fields are known to incorrectly predict secondary structural elements in IDPs and favor collapsed states.^82^ For the short P1 domain, the formation of compact states were investigated using two force-fields, Amber ff99SB-ILDN^73^ with TIP3P water model and Amber ff99SB-disp^72^ with modified TIP4P-D water model. Amber ff99SB-ILDN is a popular force field used for simulations of folded proteins which is known to correctly reproduce density of amino acid mixtures in crowded environment.^83^ Amber ff99SB-disp has optimized protein torsion angle parameters and protein-water vdW interactions to match experimental values.^72^ However, previous simulations of dipeptides using Amber ff99SB-disp failed to phase separate in direct coexistence simulations, although phase separation was observed for Amber ff99SB-ILDN.^53^

As mentioned before, in our simulation phase separation of P1 domain is observed using both force fields, although the onset of phase separation occurs at relatively higher temperature (293 K) for the Amber ff99SB-disp force field, below which the peptides remain dispersed in water (Fig 2a, *upper panel*). Above 293 K, qualitatively similar condensate formation behavior is observed between two force-fields (Fig 2a, *lower panel*). Fig 2b shows that P1 chains are more likely to become part of the largest cluster (higher clustering degree) for Amber ff99SB-ILDN force field. This may be due to the known tendency of this force field to favor compact states.^82^ Compared to Amber ff99SB-ILDN, aggregation propensity values have a larger range for Amber ff99SB-disp force field as the initial chain configuration remains dispersed with high SASA values and significant changes in SASA values are observed as the chains form the dense phase.

**Figure 2:**
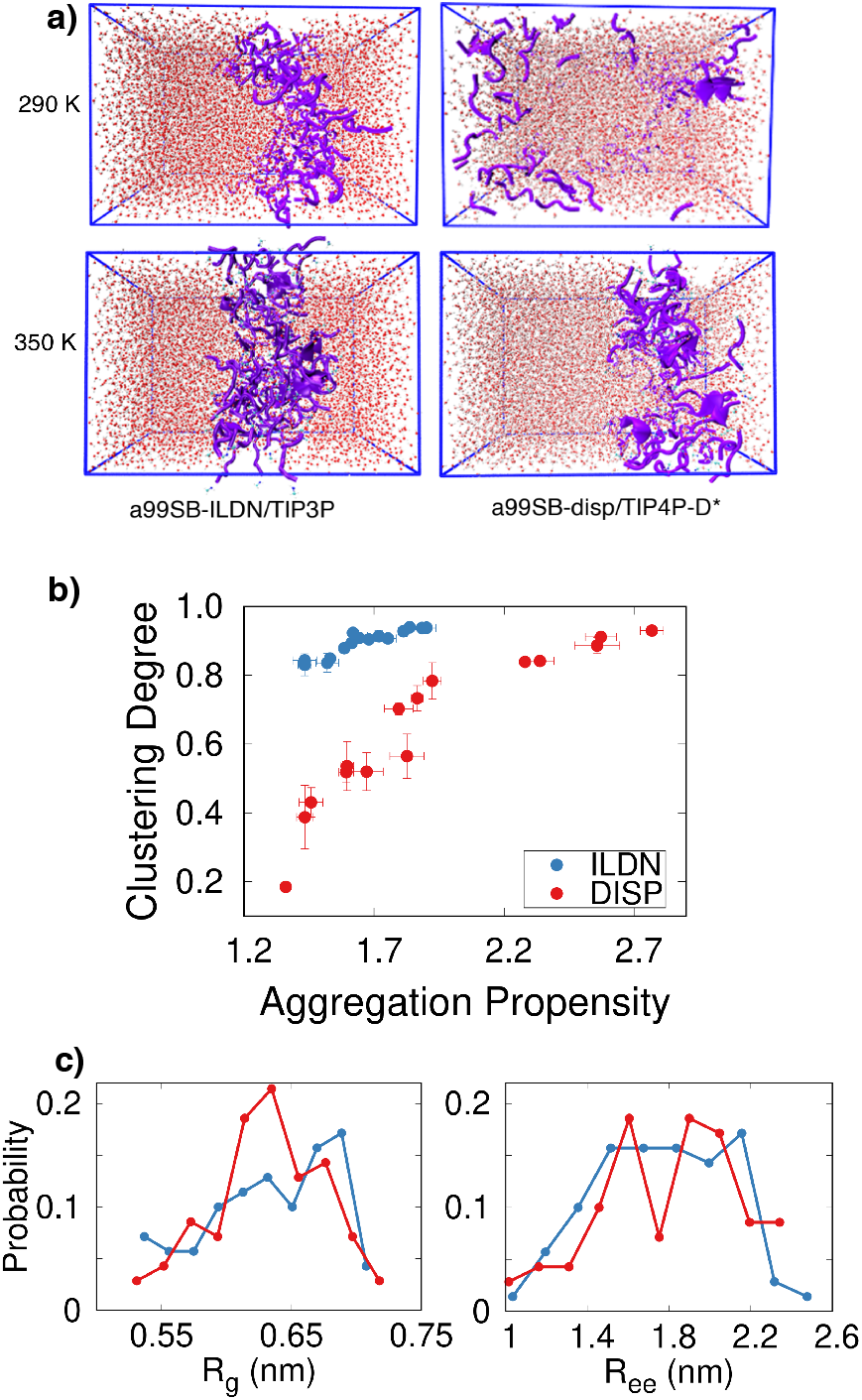
Condensate formation using different force fields. (a) Snapshot of the simulation box at 290 K and 350 K employing Amber ff99SB-ILDN force field with TIP3P water (a99SB/TIP3P) and Amber ff99SB-disp force field with modified TIP4P-D water (a99SB-disp/TIP4P-D^*^) models. (b) Projection of simulation data using two force-fields on aggregation propensity and clustering degree. (c) Distribution of time-averaged (for last 300 ns) radius of gyration *R*_*g*_ and end-to-end distance (*R*_*ee*_) at 350 K for all P1 chains using two force-fields.

To investigate P1 chain conformations in the condensed phase we obtained the distributions of the time-averaged radius of gyration (*R*_*g*_) and end-to-end distance (*R*_*ee*_) of all the chains using both the force fields (Fig. 2c). It was found that *R*_*g*_ and *R*_*ee*_ values span the same range in both the cases. The average radius of gyration exhibited by one P1 chain, in presence of other chains in the dense phase, employing Amber ff99SB-disp force field is 0.63 nm, which is the same obtained using Amber ff99SB-ILDN force field. The end-to-end distance values for disp and ILDN force fields are also similar, 1.79 and 1.76 nm, respectively.

As the results obtained using both the force-fields remain qualitatively similar, we investigate the details of condensate formation using Amber ff99SB-disp/TIP4P-D^*^ model system in the following sections. Comparison between the two force fields is presented in the ESI†, if not mentioned separately.

### 3.2 Temperature dependent phase behavior of P1

In our simulations a homogeneously mixed protein-water system separates into protein-rich and protein-dilute phases. In order to investigate the properties of the condensed phase we calculated the average density profiles of the chains along the long axis (𝒵-axis) of the simulation box at different temperatures (Fig. 3a). It was found that the densities of the condensed phases are, in general, higher at higher temperatures suggesting LCST phase behavior. To further validate our findings, simulations using Amber ff99SB-disp and modified TIP4P-D water have been repeated at 271 K, 296 K, 320 K, 335 K, 350 K and similar density profiles were obtained (data not shown).

**Figure 3:**
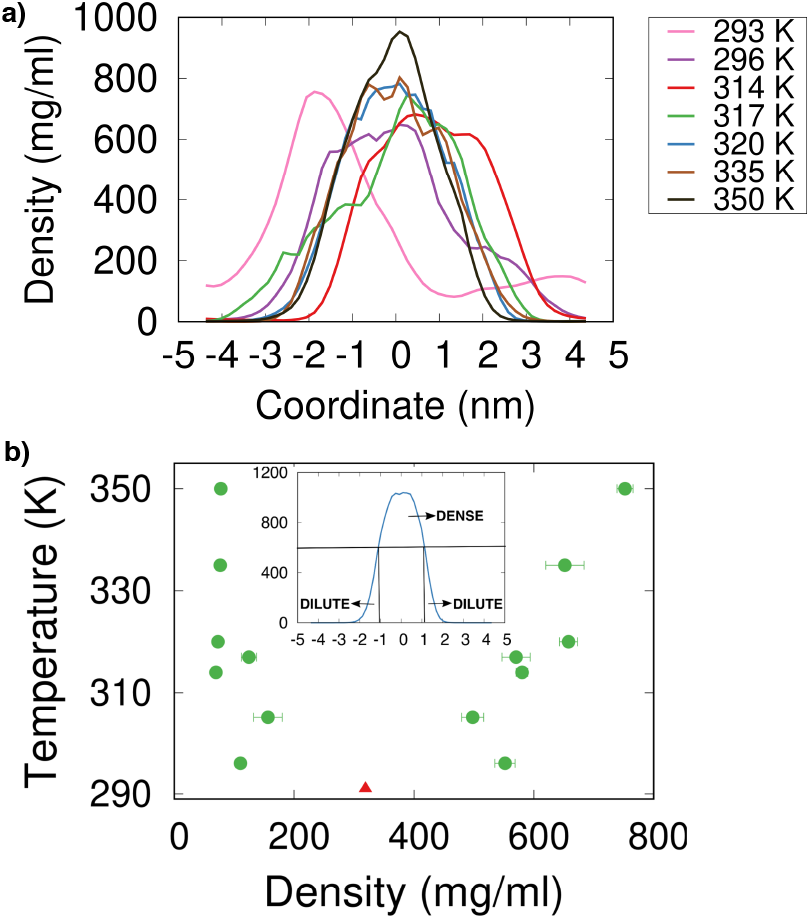
(a) P1 chain density profiles at various temperatures. (b) Temperature-density phase diagram for P1 chains. The fitted *T*_*c*_ value is shown as a red triangle. The *inset* illustrates the cut-off based method used to calculate dense and dilute phase densities.

Calculation of dilute and dense phase densities in phase coexistence simulations of IDPs remains a challenging task.^56^ In this work the dilute and dense phase densities were calculated using different cutoffs on the density profiles (see ESI Fig.S2)†, although the results remain qualitatively independent of the choice of cutoff (data not shown). Using the density profile along the long axis, the dilute and dense phases were separated by a cutoff at half-maxima of the peak value. Densities above the half-maxima were considered to be in the protein-dense phase, and all those below were interpreted to be in the dilute phase. For each of the temperatures where a coexistence was observed, the dense and dilute phase densities were estimated. The critical temperature was obtained by fitting the dense and dilute phase densities to the critical exponent equation (see Methods) with *β*=0.325. The fitting was performed for various temperature ranges. The optimum critical temperature *T*_*c*_=291 K was calculated by minimizing the error of fitting (see ESI Table S2 and S3 for calculation using two force fields)†.

In simulation, the densities of the dilute and dense phases range between 60-160 mg/ml and 500-800 mg/ml, respectively. These values are slightly higher than the experimentally determined densities of proteins in condensed phases. The density of FUS-LCD at 298 K, 150 mM NaCl, and pH 5.5 was measured to be ∼477 mg/ml.^84^ In another study the densities of a recombinant FUS-LCD in dense and dilute phases have been reported to be 15mM (320 mg/ml) and ∼60*µ*M (1 mg/ml), respectively, at room temperature and physiological salt concentration.^85^ With Amber99sb-ILDN force field parameters, the dilute and dense phase densities are 50-120 mg/ml and 600-850 mg/ml, respectively, which are also high compared to experiments.

### 3.3 Weak multi-chain contacts likely determine LCST phase behavior

Inter-chain interactions between P1 chains governed by amino acid sequences drive the formation of dense phase. In phase-separating macromolecular solutions polymers can exhibit upper critical solution temperature or lower critical solution temperature phase behavior.^18^ In order to elucidate the underlying interactions leading to the LCST phase behavior of the P1 chains, we examine the interchain contact formation among the peptides. Four temperatures, 296 K, 320 K, 335 K, and 350 K, were selected to understand the network of P1-P1 contacts at low, intermediate, and high temperatures. In simulations, contacts form between two or more P1 chains. To find out the cut-off distance for contact formation between P1 chains, we have visually identified two P1 chains that form contacts between same pairs of residues during the entire length of the simulation and obtained the average minimum distances between the C*α* atoms of different residue pairs of two interacting P1 chains. The average minimum distance is ∼1 nm at 350 K with a standard deviation of 0.04 nm. Hence, a cut-off value of 1.1 nm was chosen for contact formation between P1 chains. Since there are 7 amino acids per chain, a maximum of 7 C*α*-C*α* contacts can form between two interacting chains.

The network characteristics of contact formation between chains is obtained from the time series of the number of contacts for any pair of chains in the box. For any pair of chains the probability to form *n* or more contacts during the simulation is given by

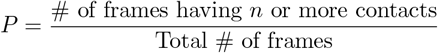

High P value indicates stable contact, whereas low P value indicates weak (unstable) contacts. We then investigate how many chain pairs have *n* or more contacts with a probability value of *x*. For example, a high probability (say, 0.8 or more) to form 6 or more contacts between two chains shows formation of stable contacts between the chains only, since there can be a maximum of 7 contacts between two chains. However, a high probability value to form 3 or more contacts indicates formation of stable contacts between multiple chains, including those formed between two chains. Thus, the cumulative increase in the number of chain pairs forming *n* or more contacts, where *n* sequentially decreases from 6 to 1, can be used as a proxy for increase in the number of multi-chain contacts.

The probability of contact formation between pairs of chains varies with temperature (see ESI Table S4, S5 and Fig. S5)†. A strong (stable) contact is defined if probability P is 0.8 and above, whereas for a weak (unstable) contact P takes values within 0.3-0.6. Fig. 4 shows the change in the number of chain pairs having *n* or more weak or strong contacts at various temperatures in the range 296-350 K. In general, with increasing temperature number of weak contacts increases, but number of strong contacts initially increases a little and then decreases slightly above 320 K. This trend is better observed in the number of multi-chain (say, 3 or more) weak contacts which increases more rapidly than the decrease in the multi-chain strong contacts as temperature goes up (Fig.4). We also calculated the number of weak contacts per chain at 296 K and 350 K (ESI Fig. S6 and S7)†, showing that the average number of weak contacts per chain at 350 K is 2.6, whereas, at 296 K it is 1.3. Hence, it is likely that increase in the number of weak multi-chain contacts at higher temperature results in the increase of density.

**Figure 4:**
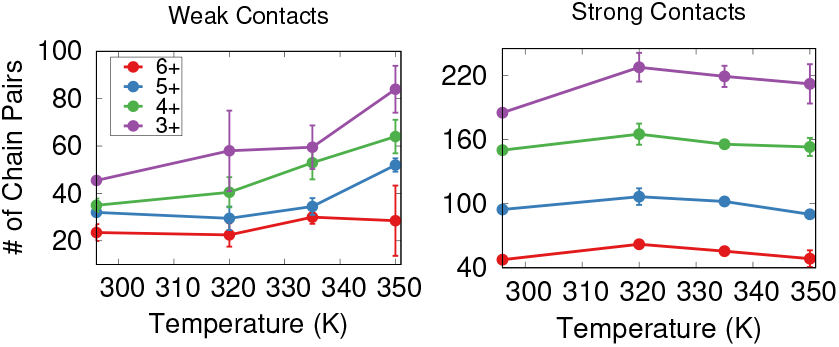
Number of chain pairs forming *n* or more weak (P value 0.3-0.6) or strong (P value 0.8 or above) contacts at different temperatures. The label *n*+ indicates *n* or more contacts.

LCST type of phase behavior of proteins is often associated with the presence of hydrophobic residues, and Glycine or Proline rich motifs.^25^ The P1 peptide is enriched in hydrophobic residues such as Valine, Leucine, and Tyrosine. To find out the driving force for P1 chain self-assembly, we have analyzed the protein-protein and protein-water hydrogen bonds. A hydrogen bond is considered when the donor and acceptor heavy atoms were at a distance of 3.5Å or lower, and the angle formed by acceptor, hydrogen atoms, and donor was 30° or less. For the final 100 ns, the hbond formation has been examined at temperatures 296 K, 320 K, 335 K and 350 K. It is found that the average number of protein-protein hbonds increases with temperature, whereas the average number of protein-water hbonds decreases (Fig. 5a), indicating that phase behavior is governed by hydrophobic interactions.

**Figure 5:**
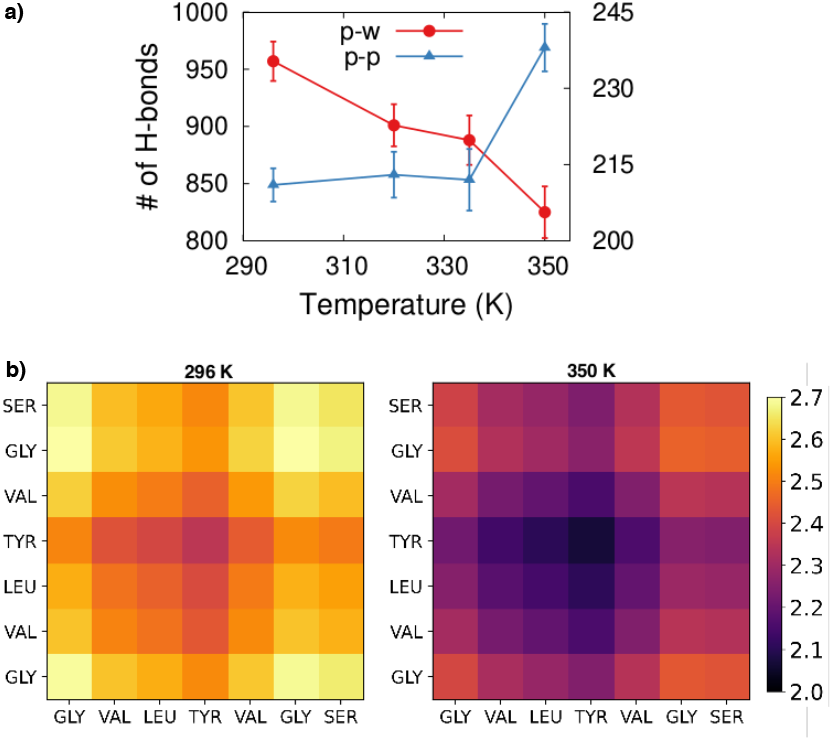
Driving force for condensate formation. Number of protein-protein (p-p) and protein-water (p-w) hydrogen bonds at varying temperatures. b)Residue-residue contact map averaged over simulation time and pair of chains at temperatures 296 and 350 K. The colorbar shows distances in nm.

Tyrosine is referred to as a sticker and can form strong *π* − *π* contacts mediating a phase separation.^86^ Highly hydrophobic residues such as Valine and Leucine can also interact with Tyrosine. To find out which amino acids form strongest contacts between P1 chains, Fig. 5b shows time-averaged residue-residue minimum distances for all pairs of residues from two chains (see ESI Fig. S7 for Amber ff99SB-ILDN force field†). At 296 K, P1 chains are observed to make contacts among Tyr-Tyr and Tyr-Leu. At a higher temperature of 350 K, it is found that P1 chains are susceptible towards making strongest contacts (say, minimum distance 2.2 nm or below) between Tyr-Tyr, Tyr-Val, Tyr-Leu. Additionally, strong contacts are also seen for Val-Leu, Val-Val and Leu-Leu. Thus, it can be inferred that the dense phase of the peptides is strongly mediated by inter-residue contacts formed by the Tyrosine residue in the middle.

## 4 Conclusions

In this work, we have performed multiple allatom MD simulations of the P1 domain at different temperatures elucidating that they phase separate above a lower critical solution temperature. The analysis reveals that hydrophobic interactions govern the phase behavior of P1 domain and weak multi-chain contacts likely result in increase in condensate density with rise in temperature. A densely connected network of P1 chains makes LLPS thermodynamically favorable and weak interactions help maintain liquid-like properties. Moreover, Tyrosine forms strongest contacts in the dense phase and is likely to play a crucial role in condensate formation. It is also seen that at any temperature the number of chain pairs forming strong contacts are larger that the chains making weak contacts (ESI Table S4 and S5)†. However, the cut-off criteria used here to define contacts between residues uses average C*α*-C*α* distances for strongly bound chain pair observed in simulations. In order to define weak contacts a larger cut-off value may be more appropriate.

At low temperatures just above the *T*_*c*_, P1 chains show linear diffusion in the condensate (data not shown). As the temperature increases, different regimes of diffusion can be observed, as has been reported before for diffusion in dense polymeric environments.^87^ It seems likely that at higher temperature, P1 chain diffusivity can be higher due to rapid formation and breaking of weak contacts, although multi-chain weak contacts help in maintaining a well-connected cluster of chains with high density. An analysis of P1 chain dynamics in the condensate goes beyond the scope of this work and will be presented in a future study.

Recent all-atom simulations of a full-length IDP (FUS protein) using various force fields investigated the influence of the force-fields and water models on the single-chain properties of the protein as well as on the stability of FUS-RNA complex.^81^ It was found that Amber ff99SB-disp with modified TIP4P-D water model provides an optimum solution for simulation of both structured and disordered peptide regions. However, the thermodynamics of phase separation and condensate properties for short peptides (here P1) showed little dependence on the choice of force-field in our simulations.

Finally, it is to be noted that although this work demonstrates formation of condensate by P1 fragments, a comprehensive all-atom phase coexistence simulation of the full-length *α*-Syn is required to elucidate the role of this fragment in the phase behavior of the entire protein. Interactions between the constituent domains of *α*-Syn likely regulates its phase separation and aggregation behavior. These different domains may have distinct phase behaviors themselves, and unraveling how the cross-talk between the domains affect the aggregate formation is a crucial step for anti-aggregation strategies. Moreover, mutation studies in the P1 region revealed that Y39A and S42A mutations affect *α*-Syn aggregation kinetics.^34^ It remains to be investigated if these mutations influence the phase behavior of the peptide as well. It is hoped that future MD studies in these directions will shed further light on the synergy between various *α*-Syn domains in regulation of its phase behavior and function.

## Acknowledgements

The authors gratefully acknowledge access to the Bioinformatics Resources and Applications Facility (BRAF), C-DAC, Pune and HPC facilities of NIT Rourkela for running the simulations.

## Notes

### Competing Interest Statement

The authors have declared no competing interest.

