## Supporting Information for "Insight into the Thermo-responsive Phase Behavior of P1 Domain of *α*-synuclein using Atomistic Simulations"

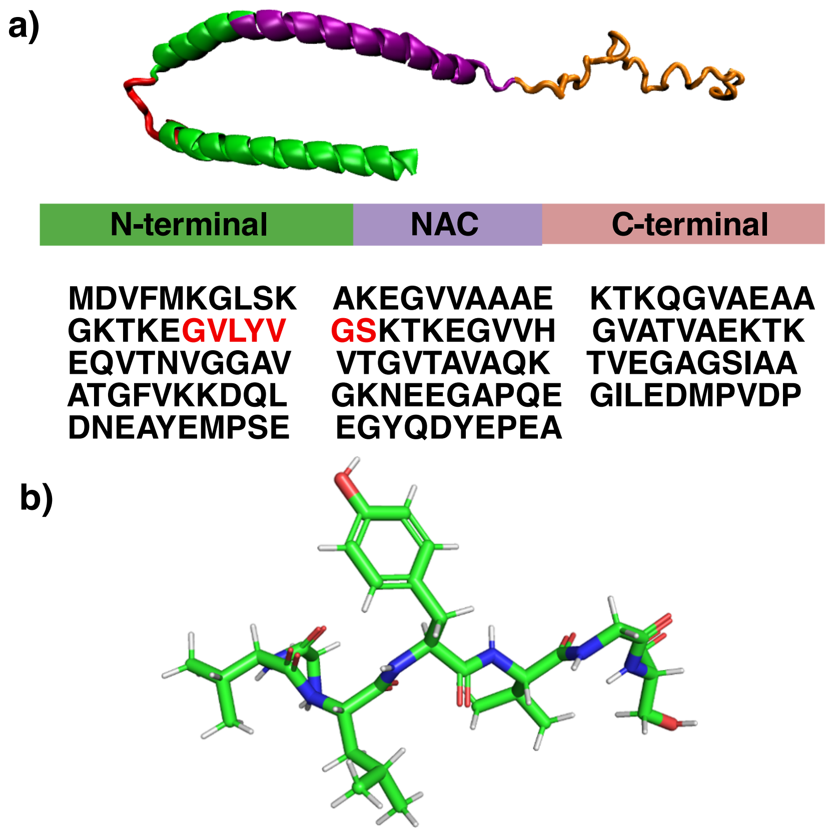


Figure S1 - (a) $\alpha$-Syn (PDB ID - 1XQ8) with the N-terminal, NAC, and C-terminal region highlighted in green, purple, and orange colours respectively, and sequence of 140 amino acid residues. (b) P1($^{36}$GVLYVGS$^{42}$) peptide structure.

**P1 System Details**

| P1 Simulation Details | | | | | |
|---|---|---|---|---|---|
| **Box Size ($nm^3$)** | **No. of protein molecules** | **No. of water molecules** | **Total no. of atoms** | **Force Field** | **Water Model** |
| 6×6×9 | 70 | 7933 | 31429 | Amberff99SB-ILDN | TIP3P |
| 6×6×9 | 70 | 8087 | 39978 | Amberff99SB-disp | TIP4P-D* |
| 6×6×12 | 70 | 11392 | 41806 | Amberff99SB-ILDN | TIP3P |

Table S1 - Simulation of P1 peptides employing various force-fields. For the system consisting of 70 P1 chains in a box of dimensions 6×6×9 $nm^3$, simulations have been conducted at temperatures 265K, 271K, 276K, 284K, 287K, 290K, 293K, 296K, 299K, 305K, 314K, 317K, 320K, 335K, 350K, and 383K using Amber ff99SB-ILDN force field with TIP3P water (a99SB/TIP3P) and Amber ff99SB-disp force field with modified TIP4P-D water (a99SB-disp/TIP4P-D*) models. The extended box ( 6×6×12 $nm^3$) simulation using Amberff99SB-ILDN force field is perfomed at 383 K only.

**Effect of box dimension on density profile of P1 chains**

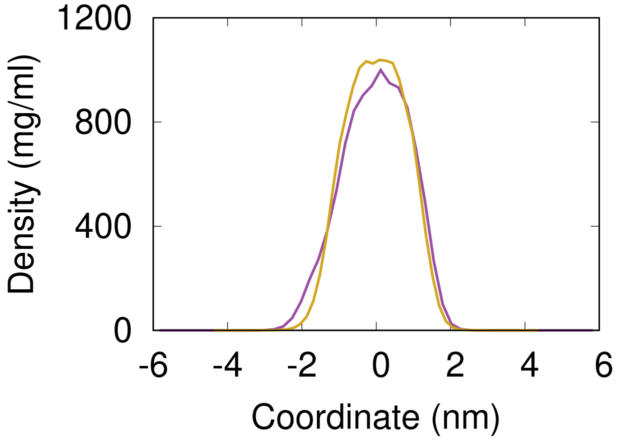


Figure S2 - Density profile of 70 P1 chains along z-dimension of the extended box (purple) and standard box (yellow) for Amber ff99SB-ILDN/TIP3P systems. In order to investigate the effect of box dimensions on the density profile of P1 chains, we simulated 70 P1 peptides in an extended box of dimensions $6\times6\times12$ $nm^3$ at a temperature of 383 K. The density behavior at a particular temperature was not found to differ greatly within a larger box.

**Calculation of dense and dilute phase densities for establishing the phase diagram.**

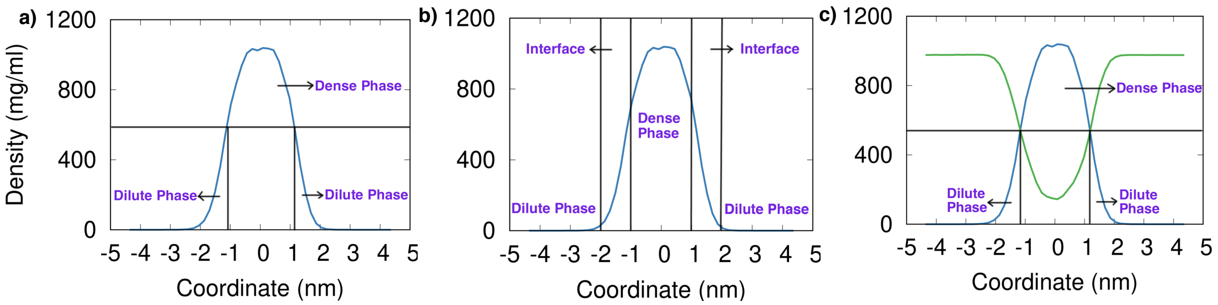


Figure S3 - Determination of dense and dilute phase densities using various approaches.

In this work, we attempted to calculate the dense and dilute phase densities by different approaches as listed below.

- We considered the half maxima of the density profile as the cut off for determining the densities. All the values lying above the half maxima of the density plot were considered to be in the dense phase, and all those below were interpreted to be in the dilute phase (Figure S2(a)). For each of the temperatures where a coexistence is observed, the dense and dilute phase densities have been approximated.

- We assumed the dense phase to lie between -1 nm $< z <$ 1 nm, and considered an interface from 1 nm $< \mid z \mid <$ 2 nm. The rest was assumed to be in the dilute phase. Then the corresponding dense and dilute protein densities were calculated (Figure S2(b)). However, the resulting phase diagram was found to overestimate the respective density values. This can be attributed to the lack of relatively flat density profiles of the peptides.

- We obtained the density profiles of the water content within the box. Thereafter, the intersection of the protein and water density profiles was chosen as the cut off for determining the dense and dilute phases (Figure S2(c)). The values obtained in this case were similar to our previous calculations using the half maxima as a cut off.

**Phase Diagram for Amber ff99SB-ILDN forcefield**

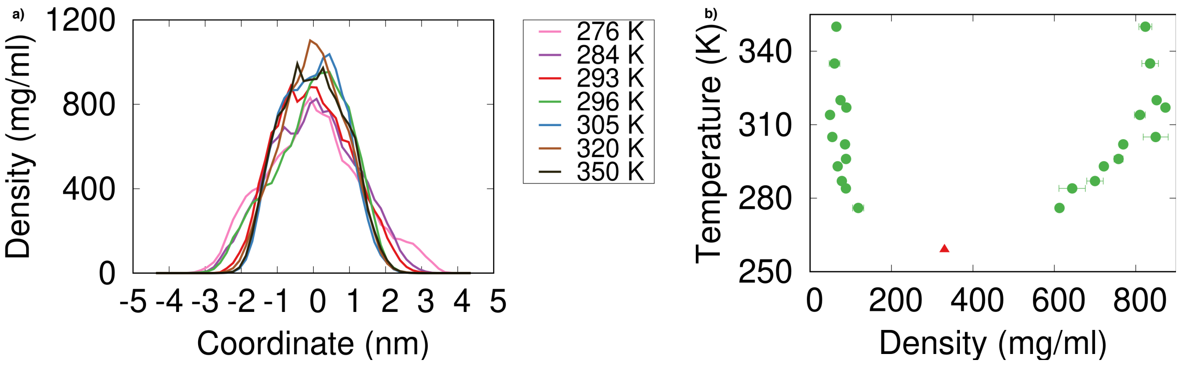


Figure S4 - (a) P1 peptide density profiles along the z-axis at different temperatures (b) Coexistence phase diagram with fitted $T_c$ value for the system built using Amber ff99SB-ILDN/TIP3P model.

To minimize the relative error of fitting, the minimum fitting temperature is set to 293 K, and the maximum fitting temperature is chosen to be 320 K. The corresponding critical temperature is estimated to be 259 K (Figure S3(b)).

We did similar calculations for the system built using Amber ff99SB-disp/TIP4P-D*. Here, the system starts to phase separate at around 293 K. For each of the temperatures where a coexistence was observed, the dense and dilute phase densities have been estimated. After considering the relative error of fitting, the minimum fitting temperature is set to 296 K, and the maximum fitting temperature is chosen to be 350 K. In this case, we obtain a critical temperature of 291 K.

**Variation of critical temperature with choice of temperature range.**

| **Tmax = 350K** | | |
|---|---|---|
| **Tmin** | **Tc** | **Error** |
| 265 | 217 | 0.06 |
| 276 | 239 | 0.06 |
| 284 | 237 | 0.04 |
| 287 | 218 | 0.04 |
| 290 | 203 | 0.04 |
| 293 | 207 | 0.04 |
| **Tmax = 320K** | | |
| **Tmin** | **Tc** | **Error** |
| 265 | 228 | 0.07 |
| 276 | 254 | 0.05 |
| 284 | 259 | 0.03 |
| 287 | 253 | 0.03 |
| 290 | 250 | 0.03 |
| 293 | 259 | 0.02 |

Table S2 - Change in critical temperature $T_c$ with varying range of fitting temperature and corresponding relative error of fitting using Amber ff99SB-ILDN/TIP3P model parameters. In each of the cases $T_{max}$ was fixed to a value and $T_c$ was estimated for varying $T_{min}$ values.

| **Tmax = 350K** | | |
|---|---|---|
| **Tmin** | **Tc** | **Error** |
| 293 | 284 | 0.20 |
| 296 | 291 | 0.17 |
| **Tmax = 320K** | | |
| **Tmin** | **Tc** | **Error** |
| 293 | 279 | 0.25 |
| 296 | 289 | 0.23 |

Table S3 - Change in critical temperature with varying range of fitting temperature and corresponding relative error of fitting using Amber ff99SB-disp/TIP4P-D* model parameters. In each of the two cases $T_{max}$ was fixed to a value and $T_c$ was estimated for varying $T_{min}$ values. $T_{min} > 296$K was not considered as it gives few data points for fitting.

**Determination of multichain contacts**

To elucidate the number contacts that each of the peptide chains make, we performed a contact analysis. The P1 peptide consists of seven amino acid residues. Thus, upon grouping the contacts made by the C$\alpha$ atoms, each peptide chain is able to make a maximum of 7 contacts. The contact probability is obtained by dividing the number of frames in which a contact is formed by the total number of frames in the final 100 ns of the simulation. The weak contacts are assumed to have a probability lying in range 0.3 to 0.6, and the strong contacts have probability in 0.8 to 1.0. Using in house Python scripts, we were able to deduce the number of chain pairs making contacts with each other. The number of contacts and the chain pairs for both the systems are showcased below.

**Amber ff99SB-disp forcefield/TIP4P-D***

| All contacts | | | | |
|---|---|---|---|---|
| **Number of Contacts** | **296 K** | **320 K** | **335 K** | **350 K** |
| 6 or more | 222±17 | 258±6 | 256±5 | 320±21 |
| 5 or more | 306±27 | 349±14 | 340±2 | 411±24 |
| 4 or more | 386±38 | 462±12 | 446±0 | 521±12 |
| 3 or more | 495±26 | 575±16 | 551±2 | 663±23 |

| Weak contacts: probability 0.3-0.6 | | | | |
|---|---|---|---|---|
| **Number of Contacts** | **296 K** | **320 K** | **335 K** | **350 K** |
| 6 or more | 23±3 | 22±4 | 30±2 | 28±14 |
| 5 or more | 32±0 | 29±4 | 34±3 | 52±2 |
| 4 or more | 35±2 | 40±6 | 53±7 | 64±7 |
| 3 or more | 45±0 | 58±16 | 59±9 | 84±9 |

| Strong contacts: probability 0.8-1.0 | | | | |
|---|---|---|---|---|
| **Number of Contacts** | **296 K** | **320 K** | **335 K** | **350 K** |
| 6 or more | 47±0 | 62±2 | 55±2 | 48±7 |
| 5 or more | 94±2 | 106±7 | 102±1 | 90±2 |
| 4 or more | 150±1 | 165±9 | 155±3 | 153±8 |
| 3 or more | 185±1 | 227±13 | 219±9 | 212±18 |

Table S4 - Number of contacts formed at four temperatures - 296K, 320K, 335K, and 350K and the corresponding average number of chain pairs for Amber ff99SB-disp forcefield and TIP4P-D* system.

**Amber ff99SB-ILDN forcefield/TIP3P**

| All contacts | | | | |
|---|---|---|---|---|
| **Number of Contacts** | **276 K** | **287 K** | **296 K** | **320 K** |
| 6 or more | 197±7 | 199±14 | 205±7 | 253±2 |
| 5 or more | 268±5 | 276±12 | 288±12 | 342±11 |
| 4 or more | 357±9 | 374±14 | 388±14 | 450±6 |
| 3 or more | 457±14 | 480±24 | 492±14 | 576±2 |

| Weak contacts: probability 0.3-0.6 | | | | |
|---|---|---|---|---|
| **Number of Contacts** | **276 K** | **287 K** | **296 K** | **320 K** |
| 6 or more | 18±3 | 18±7 | 21±2 | 29±2 |
| 5 or more | 24±5 | 21±5 | 35±5 | 32±0 |
| 4 or more | 27±4 | 33±0 | 26±14 | 64±2 |
| 3 or more | 37±0 | 33±2 | 29±4 | 62±2 |

| Strong contacts: probability 0.8-1.0 | | | | |
|---|---|---|---|---|
| **Number of Contacts** | **276 K** | **287 K** | **296 K** | **320 K** |
| 6 or more | 69±2 | 61±2 | 61±1 | 65±0 |
| 5 or more | 112±5 | 114±3 | 115±3 | 120±0 |
| 4 or more | 181±1 | 182±2 | 185±10 | 186±4 |
| 3 or more | 243±9 | 259±9 | 263±11 | 264±5 |

Table S5 - Number of contacts formed at four temperatures - 276K, 287K, 296K, and 320K and the corresponding average number of chain pairs for Amber ff99SB-ILDN forcefield and TIP3P water model system. These temperatures were chosen as the condensate density increases with increasing temperature within the range 265K - 320K (Fig. S4 (b)).

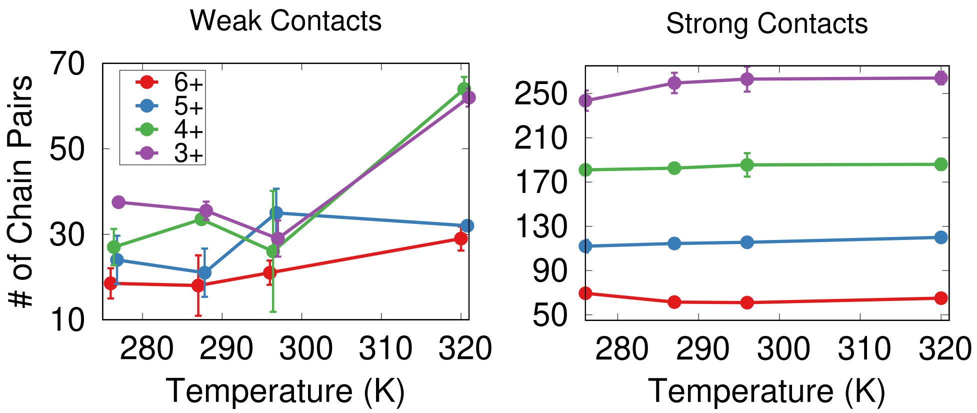


Figure S5 - Number of chain pairs forming weak and strong contacts at different temperatures for the Amber ff99SB-ILDN/TIP3P system.

**Contacts formed by each P1 chain**

- **Amber ff99SB-ILDN forcefield/TIP3P**

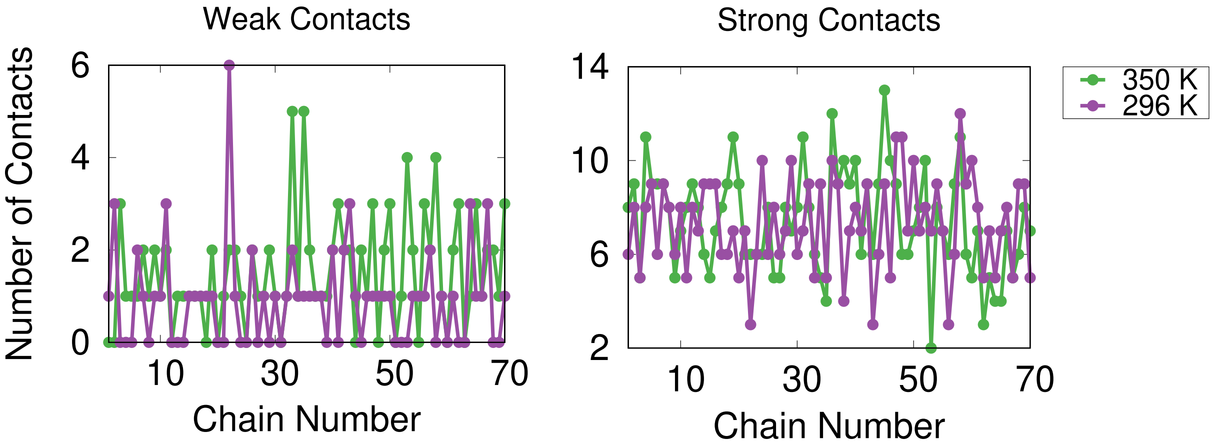


Figure S6 - Number of contacts (averaged over last 100 ns simulation) formed by each of the 70 P1 chains for Amber ff99SB-ILDN forcefield/TIP3P at 296 K and 350 K.

- **Amber ff99SB-disp forcefield/TIP4P-D***

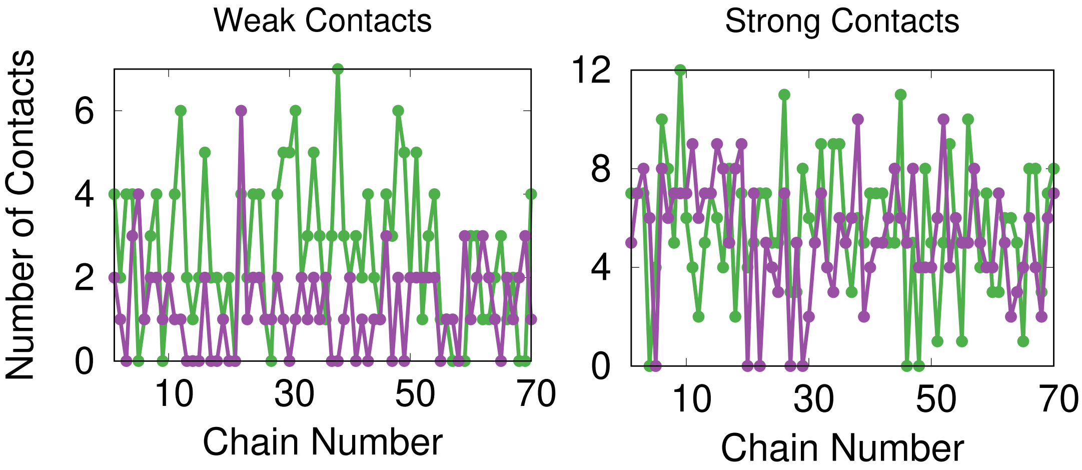


Figure S7 - Number of contacts (averaged over last 100 ns simulation) formed by each of the 70 P1 chains for Amber ff99SB-disp forcefield/TIP4P-D* at 296 K and 350 K.

**Residue level distance map**

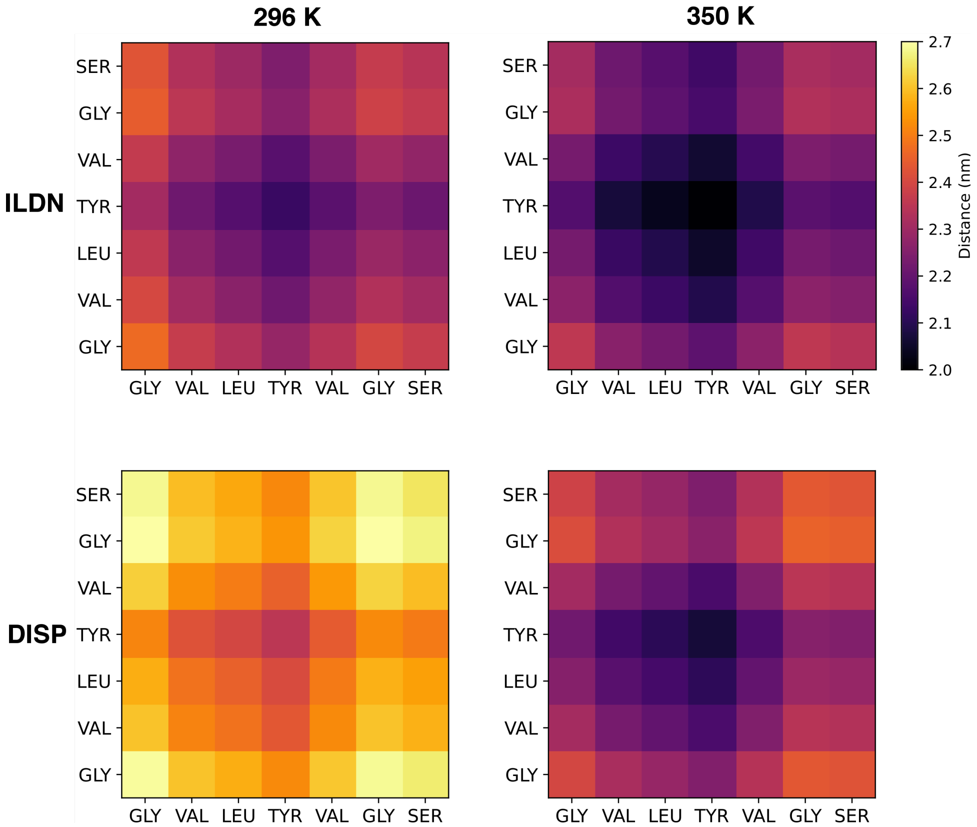


Figure S8 - Comparison of residue level average minimum distances for both force fields at 296 K, and 350 K.
